# Variation in expression of the HECT E3 ligase *UPL3* modulates LEC2 levels, gene expression, seed size and crop yields in *Brassica napus*

**DOI:** 10.1101/334581

**Authors:** Charlotte Miller, Rachel Wells, Neil McKenzie, Martin Trick, Joshua Ball, Abdelhak Fatihi, Bertrand Debreuq, Thierry Chardot, Loic Lepiniec, Michael W Bevan

**Affiliations:** Department of Cell and Developmental Biology, John Innes Centre, Norwich UK; Institut Jean-Pierre Bourgin, INRA, AgroParisTech, CNRS, Université Paris-Saclay, INRA Versailles, route de Saint-Cyr, FR

**Author notes:** **Corresponding author** Michael W. Bevan. Orcid ID 0000-0001-8264-2354, Cell and Developmental Biology Dept, John Innes Centre, Norwich Research Park, Norwich NR4 7UH, UK.

## Abstract

Identifying genetic variation that increases crop yields is a primary objective in plant breeding. We have used association analyses of *Brassica napus* (oilseed rape/canola) accessions to identify variation in the expression of a HECT E3 ligase gene, *BnaUPL3.C03,* that influences seed size and final yield. We establish a mechanism in which UPL3 mediates the proteasomal degradation of LEC2, a master regulator of seed maturation. Reduced *UPL3* expression increases LEC2 protein levels and prolongs expression of lipid biosynthetic genes and seed maturation. Natural variation in *BnaUPL3.C03* expression has not yet been exploited in current *Brassica napus* breeding lines and can therefore be used as a new approach to maximize future yields in this important oil crop.

## Introduction

In major oil-producing crops, such as *Brassica napus* (oilseed rape), the composition of seed storage lipids has been optimized for different end uses, from human nutrition to industrial applications, by identifying allelic variation in biosynthetic enzymes and pathways(Napier and Graham, 2010). For example, elite oilseed rape varieties now have greatly enhanced nutritional value, with high linoleic acid, reduced erucic acid and optimal linoleic to linolenic acid ratios. However, increasing overall production of storage lipids to meet projected future demands for both food and industrial uses remains a key objective for achieving food security and sustainable industrial production.

An interacting network of transcription factors establishes and maintains embryo development and promotes the accumulation of seed storage lipid and protein(Fatihi et al., 2016; Boulard et al., 2017). Loss-of-function mutations in four conserved regulatory genes lead to curtailed seed maturation, loss of dormancy, and ectopic vegetative development. *FUSCA3* (*FUS3*), *ABSCISIC ACID INSENSITIVE 3* (*ABI3*) and *LEAFY COTYLEDON 2* (*LEC2*) encode AFL-B3-family transcription factors, while *LEAFY COTYLEDON 1* (*LEC1*) encodes an NFY-Y CCAAT binding transcription factor. These “*LAFL*” genes(Santos-Mendoza et al., 2008a; Roscoe et al., 2015) induce seed maturation and inhibit germination, and their expression is down-regulated at the initiation of seed dormancy and desiccation tolerance. *LEC2* and *FUS3* expression is repressed by miRNA-mediated mechanisms during early embryogenesis(Nodine and Bartel, 2010; Willmann et al., 2011) to ensure the correct timing of storage reserve accumulation. At later stages of seed development, the B3-domain protein VAL3 recruits Histone Deacetylase19 (HDAC19) to the promoters of LAFL genes and silences their expression by altering levels of histone methylation and acetylation (Zhou et al., 2013). The stability of LAFL proteins is also controlled during seed maturation and dormancy. ABI3-interacting protein 2 (AIP2) is an E3 ligase that ubiquitylates both ABI3(Zhang et al., 2005) and FUS3 (Duong et al., 2017), suggesting that regulation of LAFL protein levels has an important role in seed maturation and the transition to dormancy. The SNF kinase AKIN10 phosphorylates and stabilizes FUS3 (Chan et al., 2017; Tsai and Gazzarrini, 2012) and WRI1 (Zhai et al., 2017), a transcription factor regulated by LEC2 that promotes expression of glycolytic and lipid biosynthesis genes. Improved understanding of the control of these important seed maturation regulators may provide new understanding for optimizing seed yields.

An important strategy in crop improvement aims to identify new sources of genetic variation from diverse germplasm resources for increasing crop productivity(Bevan et al., 2017). Genome-Wide Association Studies (GWAS) are increasingly used for identifying variation associated with traits in crops and their wild relatives. For example, associations between sequence variation, gene expression levels and oil composition have been used to identify genetic variation in known genes conferring oil quality traits in oilseed rape (Harper et al., 2012; Lu et al., 2017). GWAS also has potential for the discovery of new gene functions, and, when utilized fully, can lead to a deeper understanding of mechanisms underlying complex traits such as crop yield. Here we use Associative Transcriptomics in *Brassica napus* to identify genetic variation in the regulation of *BnaUPL3.C03,* encoding an ortholog of the HECT E3 ubiquitin ligase *UPL3 (*Downes et al., 2003; El Refy et al., 2003), that is associated with increased seed size and field yields. We establish a mechanism in which reduced expression of *BnaUPL3.C03* maintains higher levels of LEC2 protein during seed maturation by reduced UPL3-mediated LEC2 ubiquitylation, leading to increased seed lipid levels and overall increased seed yields. Analysis elite oilseed rape varieties shows that variation in the expression of *BnaUPL3.C03* has not yet been exploited in breeding programmes and thus can be used to increase crop yields.

## Results

### Associative Transcriptomics identifies a negative correlation between *BnaUPL3.C03* expression and seed weight per pod in *B. napus*

A panel of 94 *B. napus* oilseed rape accessions with high genetic diversity (Supplementary Table 1), for which leaf transcriptome data from each accession was mapped to a sequenced reference genome(Harper et al., 2012), was screened for yield-related phenotypic variation. High levels of trait variation were observed. Trait associations with both sequence variation, in the form of hemi-SNPs in the polyploid genome of *B. napus*, and gene expression levels, assessed as Reads Per Kilobase per Million (RPKM) mapped reads in a Gene Expression Marker analysis (GEM), were then calculated. This identified significant associations between variation in seed weight per pod (SWPP) and SNP variation in homoeologous regions of linkage groups A08 and C03 (Figure 1A and Supplementary Figure 1). SWPP phenotype data displayed a normal distribution appropriate for Mixed Linear Model analyses (Figure 1B).

**Figure 1.**
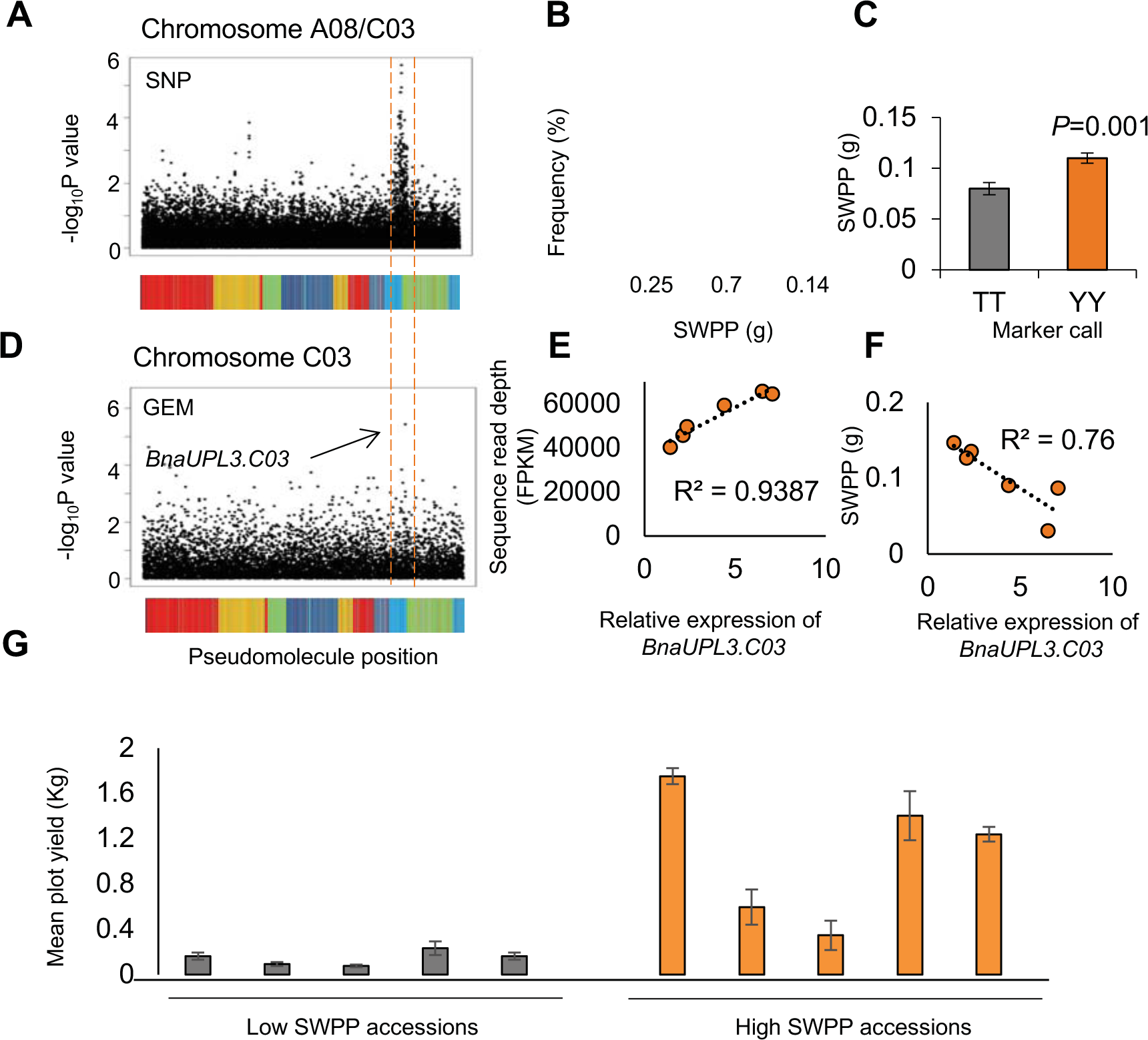
Association of variation in Seed Weight Per Pod (SWPP) with SNPs and differential expression of *BnaUPL3.C03* in a panel of 94 *Brassica napus* accessions. **A.** Associations between transcriptome single-nucleotide polymorphism (SNP) on chromosomes A08/C03 and SWPP. The dotted red lines outline a peak of SNP associations. Markers are plotted in pseudomolecule order, and associations as −log10 *P* values. The coloured regions under the pseudochromosome represent the regions of sequence similarity to *A. thaliana* chromosomes, as described previously (Harper et al. 2012). **B.** Normal distribution of the SWPP trait in the set of *B. napus* accessions. **C.** Segregation of SWPP trait means with the most highly associating marker (JCVI_5587:125) show a marker effect of ~20%. Data are given as mean ± SE. *P* values were determined by Student’s *t*-test. **D.** Gene Expression Marker (GEM) differential expression of a single C genome assigned unigene C_EX097784 on chromosome C03 is associated with SWPP variation. This unigene is an ortholog of Arabidopsis *UBIQUITIN PROTEIN LIGASE 3* (*UPL3*), and is termed *BnaUPL3*.C03. The coloured regions under the pseudochromosome represent the regions of sequence similarity to *A. thaliana* chromosomes, as described previously (Harper et al. 2012). **E.** GEM expression (FPKM) levels in six *B. napus* GWAS accessions (Supplementary Table 2) exhibiting maximal variation in C-EX097784 expression were compared to q-RT-PCR measurements of *BnaUPL3.C03* expression in seedlings, expressed relative to *BnaUBC10*. **F.** Correlation of *BnaUPL3.C03* transcript abundance in seedlings, measured using q-RT-PCR in six *B. napus* GWAS accession exhibiting maximal variation, with variation in SWPP. **G.** A subset of ten GWAS accessions with maximal variation in SWPP were grown in replicated plots in field conditions between March-Aug 2016 and mean plot yields measured after combining. Plot yields are shown as mean ± SE.

Assessment of phenotypic variation segregating with alleles for the most significant SNP marker, JCVI_5587:125, revealed that accessions inheriting “T” at this locus had SWPP values ~20% lower than those accessions inheriting the hemi-SNP genotype, “Y” (C/T) (Figure 1C). GEM analyses showed significant association of SWPP with varying expression of a single C genome-assigned unigene within this region, C_EX097784. (Figure 1D and Supplementary Figure 1). This unigene corresponded to an ortholog of the Arabidopsis *UBIQUITIN PROTEIN LIGASE 3* (*UPL3*) encoding a HECT E3 ligase (Downes et al., 2003; El Refy et al., 2003).

Two *UPL3* homologs in *B. napus, BnaUPL3.C03* and *BnaUPL3.A08*, were identified based on protein sequence similarity and conserved synteny between Arabidopsis, *B. rapa* and *B. oleracea.* Supplementary Figure 2 illustrates the high protein sequence similarity between the single Arabidopsis and two *B.napus UPL3* orthologs. Associative Transcriptomics analyses identified significant differential expression of the *BnaUPL3.C03* gene between GWAS accessions displaying high variation in SWPP (Figure 1D and Supplementary Figure 1). Gene-specific qRT-PCR analyses confirmed this differential expression at the *BnaUPL3.C03* locus in seedlings of six lines selected for high-or low-SWPP (Figure 1E). Correlating *BnaUPL3.C03* expression levels with SWPP revealed a negative relationship, with accessions displaying low *BnaUPL3.C03* exhibiting high SWPP (Figure 1F). qRT-PCR analyses of *BnaUPL3.A08* expression in the set of six lines with high-or low-SWPP showed no significant differences in expression levels and variation in SWPP (Supplementary Figure 3). This is consistent with the absence of an association between transcript levels at the *BnaUPL3.A08* locus and SWPP determined by Associative Transcriptomics analyses (Supplementary Figure 1).

**Figure 2.**
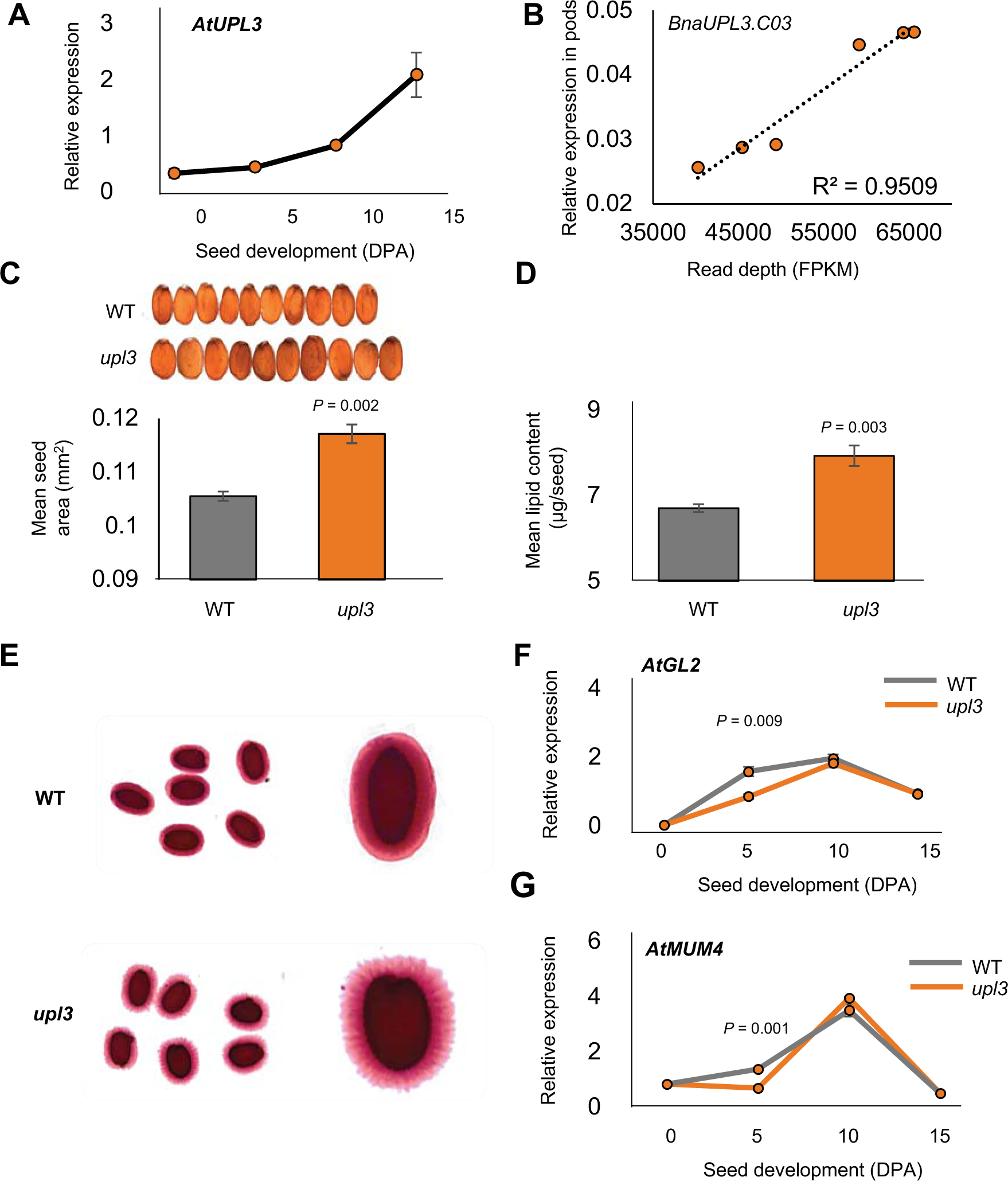
A loss-of-function mutation in Arabidopsis *UPL3* has increased seed size and altered seed storage and seed coat phenotypes. **A.** Arabidopsis *UPL3* expression, measured by q-RT-PCR, during seed development. Expression levels are relative to *EF1ALPHA* gene expression. DPA is Days Post Anthesis. Data are given as mean ± SE (n=3 per genotype). **B.** *Brassica napus BnaUPL3.C03* expression measured in maturing pods (45 DPA) across the subset of six GWAS accessions shown in Figure 1e using q-RT-PCR compared to GEM expression measured in leaves. The *BnaACTIN2* gene was used as an internal q-RT-PCR control. (n=3 per genotype). **C.** Areas of Arabidopsis seeds from WT and the *upl3* mutant. Data are given as mean ± SE (means calculated across 6 biological replicates per genotype (100 seeds per replicate) for each genotype)). *P* values were determined by Student’s *t*-test. **D.** Lipid content of Arabidopsis seeds from WT and the *upl3* mutant. Data given as means ± SE. *P* values were determined by Student’s *t*-test. **E.** Mucilage extrusion of imbibed Arabidopsis seeds from WT and the *upl3* mutant, visualised by Ruthenium Red staining. **F.** Expression profile of *GL2* in developing seeds of WT and *upl3* mutant Arabidopsis using q-RT-PCR. RNA from whole siliques was used. Expression levels are relative to *EF1ALPHA* gene expression. Data are given as means ± SE (n=3). *P* values were determined by Student’s *t*-test. **G.** Expression profile of *MUM4* in developing seeds of WT and *upl3* mutant Arabidopsis using q-RT-PCR. RNA from whole siliques was used. Expression levels are relative to *EF1ALPHA* gene expression. Data are given as means ± SE (n=3). *P* values were determined by Student’s *t*-test.

**Figure 3.**
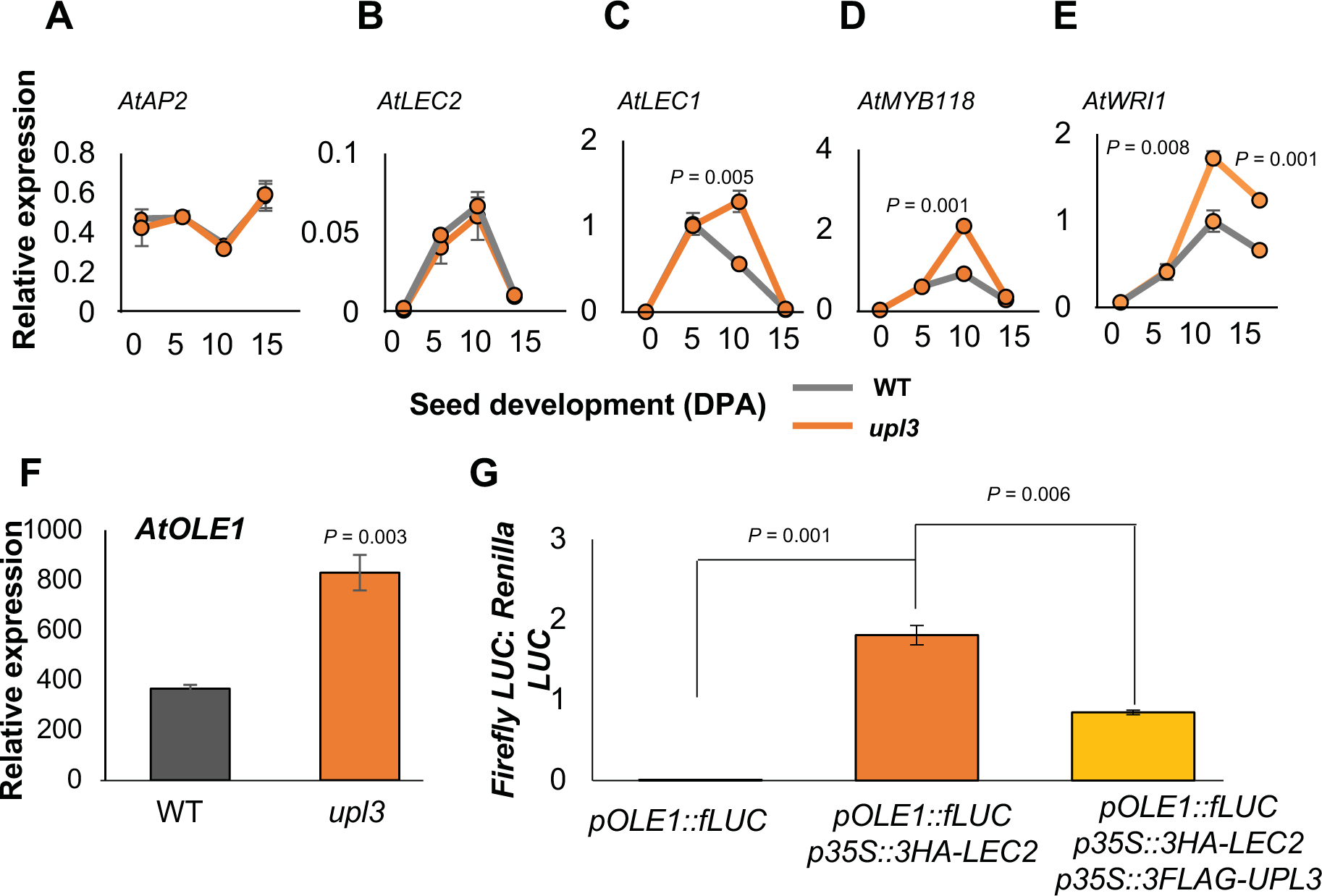
*UPL3* disrupts LEC2-mediated gene expression. q-RT-PCR was used to measure gene expression during seed development in WT and *upl3* Arabidopsis mutant plants. RNA from whole siliques was used. Expression levels are relative to *EF1ALPHA* gene expression. Data are given as means ± SE (n=3). *P* values were determined by Student’s *t*-test. **A,** *AP2* expression; **B**, *LEC2* expression; **C**, *LEC1* expression; **D**, *MYB118* expression; **E**, *WRI1* expression; **F.** q-RT-PCR measurement of *OLE1* in 10 DPA WT and *upl3* mutant Arabidopsis plants. RNA from whole siliques was used. Expression levels are relative to *EF1ALPHA* gene expression. Data are given as means ± SE (n=3). *P* values were determined by Student’s *t*-test. **G.** Transient expression of an *OLE1* promoter::*Firefly Luciferase (fLUC)* reporter gene in Arabidopsis *upl3* mutant leaf protoplasts. *35S::Renilla LUC* vector was co-transfected in all treatments as a transfection control, and the ratio of Firefly/Renilla LUC activity was used to determine *OLE1::fLUC* gene expression levels. 35S::3HA-LEC2 and 35S::3FLAG-UPL3 were co-transfected as shown. Data are presented as means ± SE. (n=3). *P* values were determined by Student’s *t*-test.

A subset of 10 GWAS accessions (Supplementary Table 2) exhibiting both differential expression of *BnaUPL3.C03* and high variation in SWPP were grown under field conditions in a replicated yield trial. Mean plot yields across accessions showed a positive correlation between SWPP values and plot yields of the accessions (R^2^=0.49) (Figure 1G), showing that SWPP is an important measure of seed yield under field conditions.

Previous studies in Arabidopsis, a close relative of *B. napus,* identified roles for UPL3 in mediating the proteasomal degradation of GLABROUS 3 (GL3) and ENHANCER OF GLABROUS 3 (EGL3), both known regulators of trichome morphogenesis. Enhanced GL3/EGL3 protein levels in a *upl3* mutant altered leaf trichome morphogenesis(Downes et al., 2003; El Refy et al., 2003; Patra et al., 2013). Assessment of leaf hairs across a subset of *B. napus* GWAS accessions with maximal variation in *BnaUPL3.C03* expression revealed segregation of a trichome phenotype (Supplementary Figure 4), suggesting differential *UPL3* expression also influences trichome morphogenesis in oilseed rape. Although *AtUPL3* appears to have a conserved role in trichome morphogenesis, there has been no evidence that *AtUPL3* has a role in seed development.

**Figure 4.**
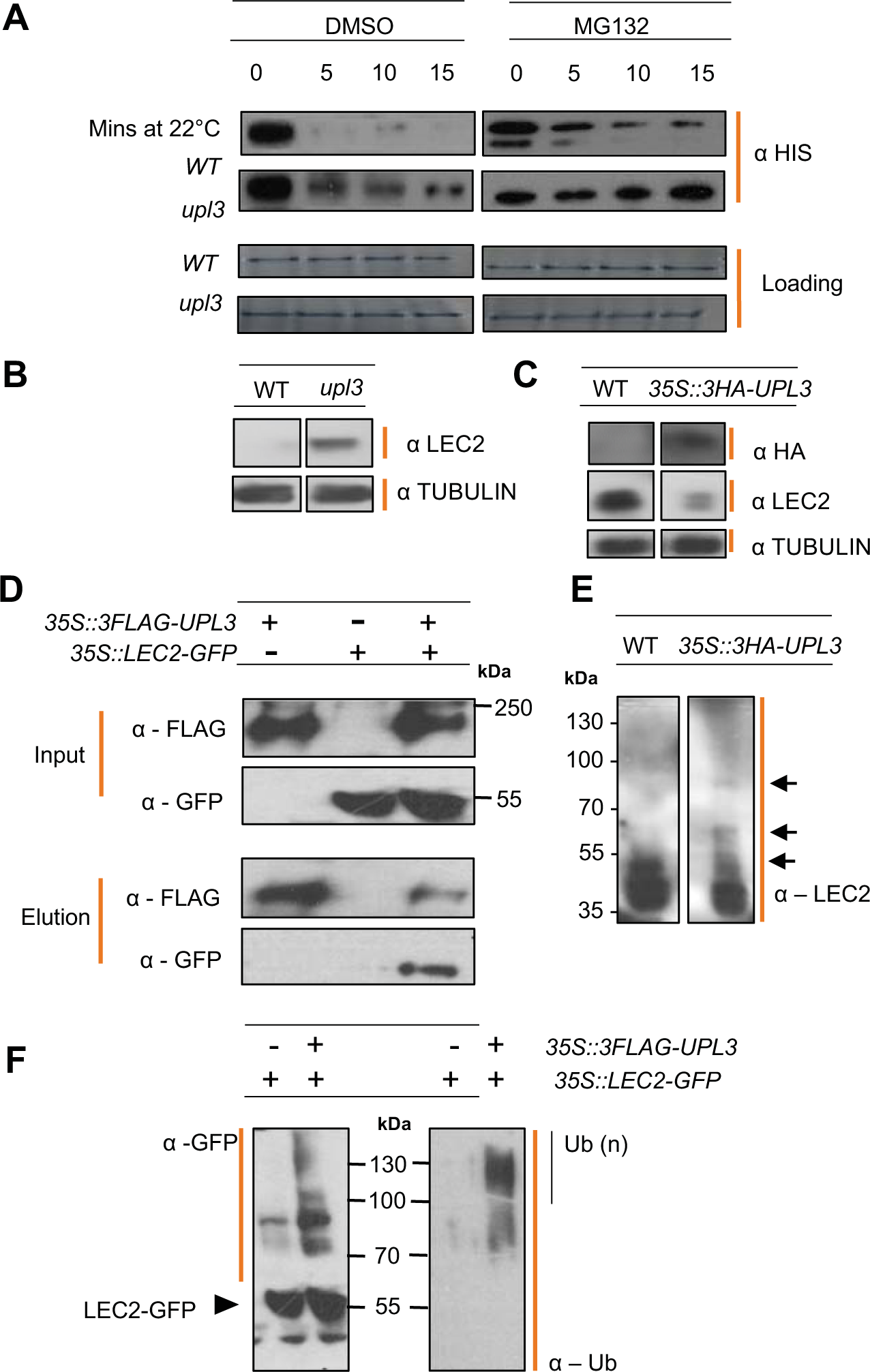
UPL3 mediates ubiquitylation and proteosomal degradation of LEC2 in Arabidopsis. **A.** Western blots of 4-20% SDS-PAGE gels probed with anti-HIS antibodies (top four panels). Purified LEC2-HIS expressed in *E. coli* was incubated at 22°C for the times indicated in total protein extracts from 10-15 DPA WT or *upl3* mutant siliques, with and without 200 μM MG132. The lower four panels are the same volume of sample taken at each timepoint electrophoresed on 4-20% SDS-PAGE gels and stained with Colloidal Coomassie Blue. **B.** Western blots of protein samples from pooled 10 and 15 DPA siliques of WT or *upl3* mutant plants electrophoresed on 4-20% SDS-PAGE gels and probed with anti-LEC2 or anti-tubulin antibodies. Tubulin levels are for comparison. **C.** Western blots of protein samples from pooled 10 and 15 DPA siliques of WT or transgenic *35S::3HA-UPL3* over-expressor plants electrophoresed on 4-20% SDS-PAGE gels and probed with anti-HA (top panels) anti-LEC2 (central panels) or anti-tubulin (lower panels) antibodies. Tubulin levels are for comparison. **D.** Western blots of protein samples from *35S::LEC2-GFP* and *35S::3FLAG-UPL3* vectors transiently expressed in *Nicotiana benthamiana* leaves and probed with anti-GFP and anti-FLAG antibodies. The upper panels show input levels of 3FLAG-UPL3 and LEC2-GFP. The lower panels show proteins purified on FLAG magnetic beads to assess co-immunoprecipitation of proteins. **E.** Western blots of protein samples shown in panel **c** from pooled 10 and 15 DPA siliques of WT or transgenic *35S::3HA-UPL3* over-expressor plants electrophoresed on 4-20% SDS-PAGE gels and probed with anti-LEC2. A longer exposure reveals higher MW forms of LEC2 protein that are indicated in the *35S::3HA-UPL3* sample. **F.** Western blots of protein samples from *35S::LEC2-GFP* and *35S::UPL3-FLAG* vectors transiently expressed in *Nicotiana benthamiana* leaves used in panel **d**. Proteins were probed with anti-GFP antibodies (left panel) and anti-ubiquitin antibodies (right panel). Higher MW forms of purified LEC2-GFP are ubiquitylated.

In Arabidopsis, *UPL3* transcript levels increased steadily during seed development, with highest expression levels observed during the seed maturation phase (Figure 2A), suggesting a potential role for *UPL3* in seed maturation. *BnaUPL3.C03* is differentially expressed in seedlings of *B. napus* accessions displaying high variation in SWPP (Figure 1F), therefore variation of *BnaUPL3.C03* expression was measured using q-RT-PCR in developing pods at 45 Days Post Anthesis (DPA) in six accessions with low- and high-SWPP. Figure 2B confirms that variation in *BnaUPL3.C03* expression in pods was tightly correlated with that in leaves measured using RNAseq.

### Arabidopsis and *B. rapa* mutants lacking *UPL3* function exhibit increased seed size

The potential influence of *AtUPL3* in seed formation was assessed using an Arabidopsis T-DNA insertion line (Salk_015534) with complete loss of *AtUPL3* expression (data not shown). *upl3* mutant seeds were approximately 10% larger (Figure 2c) and displayed a 12% increase in seed lipid content (Figure 2D) relative to seeds of WT plants. Analysis of seed lipid composition revealed no significant changes in fatty acid composition (Supplementary Table 3). A potential loss-of-function stop codon in the single *B. rapa UPL3 gene* was identified using a TILLING (Targeting Induced Local Lesions IN Genomes) resource for *B. rapa* (Stephenson et al., 2010). Supplementary Figure 5 shows that a line homozygous for the *UPL3* mutation also had larger seeds compared to a line that segregated the *UPL3* mutation. These results established a potential role for *UPL3* in regulating seed size in Arabidopsis and Brassica.

**Figure 5.**
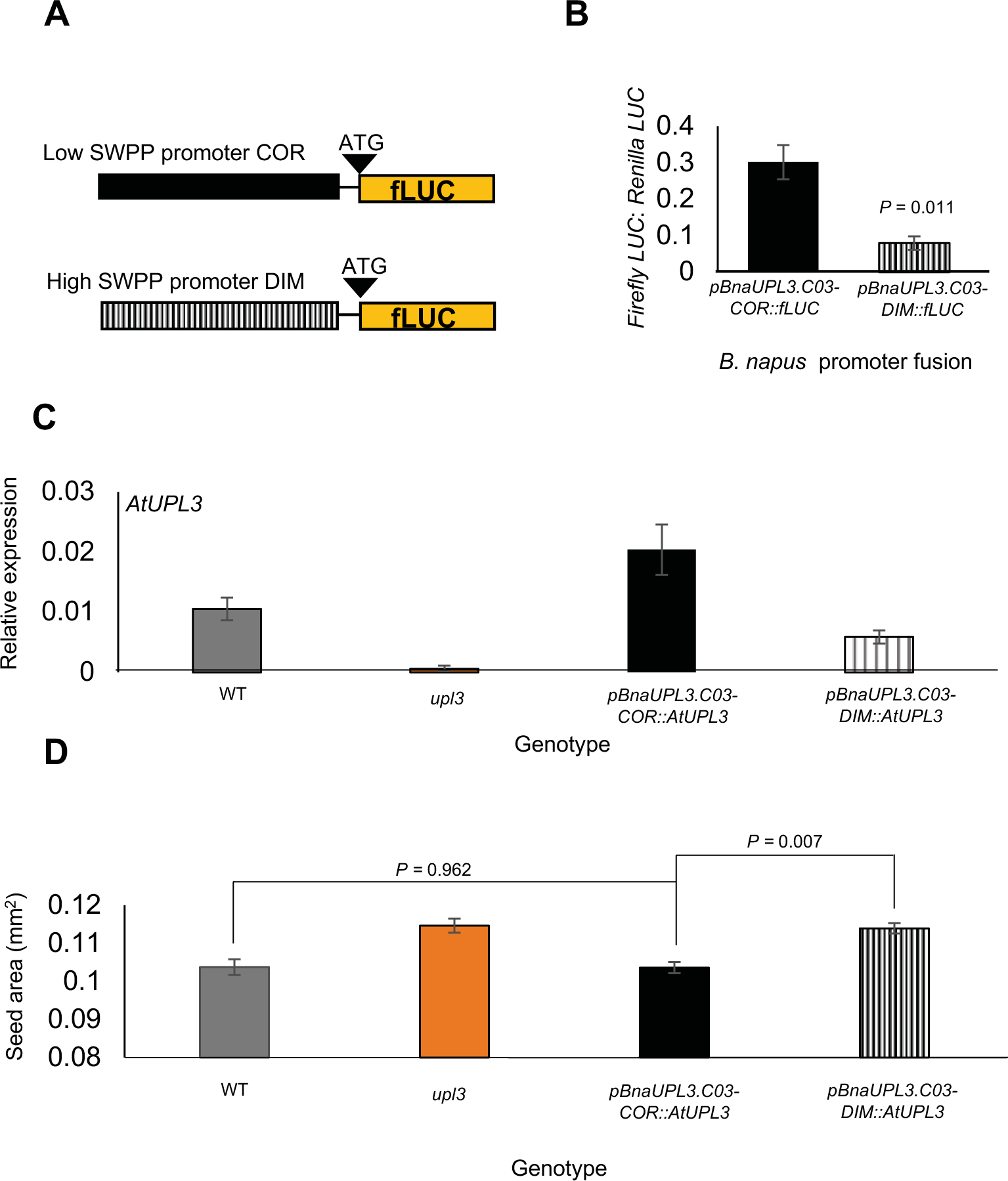
Variation in *BnaUPL3.C03* promoter activities from high- and low-SWPP B. *napus* accessions is sufficient to generate variation in final seed yield. **A.** The diagram shows fusions of 2 kb *BnaUPL3.C03* promoters from the *B. napus* lines Dimension (DIM) (High SWPP) and Coriander (COR) (Low SWPP) to the Luciferase coding region. The diagrams are not to scale. **B.** The DIM and COR promoters described in panel A were fused with a Firefly Luciferase reporter gene and transfected into Arabidopsis *upl3* mutant protoplasts. The activities of each promoter are shown relative to co-expressed *35S::Renilla* luciferase. Data are given as means ± SE (n=3). *P* values were determined by Student’s *t*-test. **C.***BnaUPL3.C03* promoters from the *B. napus* lines Dimension (DIM) and Coriander (COR) were fused to the coding region of Arabidopsis *UPL3* and transformed into *upl3* mutant Arabidopsis. q-RT-PCR of At*UPL3* expression in wild-type and and transgenic Arabidopsis lines were measured and are shown relative to the *AtEIF1ALPHA* gene. Data are given as means ± SE (at least 3 biological replicates per genotype). *P* values were determined by Student’s *t*-test. **D.** Seed area was quantified in WT, *upl3* mutant, *COR::AtUPL3* and *DIM::AtUPL3* transgenic lines. Data shown are means ± SE based on seed area measurements across at least 100 seeds per genotype and with at least 3 biological replicates per genotype. *P* values were determined by Student’s *t*-test.

### *UPL3* indirectly influences seed mucilage biosynthesis *via GL2*

Arabidopsis *upl3* mutant seeds exhibited altered mucilage extrusion upon imbibition (Figure 2E). UPL3-mediates the proteasomal degradation of the bHLH transcription factors GL3 and EGL3 (Patra et al., 2013). These proteins function as part of a complex to regulate the expression of *GLABROUS 2 (GL2)*, encoding a homeodomain transcription factor that activates expression of *MUCILAGE-MODIFIED 4* (*MUM4),* encoding a mucilage biosynthetic enzyme (Shi et al., 2012). This EGL3/GL3 complex activates GL2 expression in leaves, but represses GL2 expression in seeds, depending on the type of MYB transcription factor in the complex (Song et al., 2011). The single repeat MYB *GmMYB73*, a possible homolog of Arabidopsis *TRY* and *CPC*, repressed GL2 expression in seeds and interacted with EGL3 and GL3 (Liu et al., 2014). Thus, reduced UPL3-mediated destabilization of EGL3 and GL3 in the *upl3* mutant may elevate GL3 and EGL3 levels, leading to increased repression of GL2 expression. A significant reduction in the expression of both *GL2* (Figure 2f) and *MUM4* (Figure 2G) was observed at 5 DPA in developing seeds of the *upl3* mutant, and may explain the altered mucilage extrusion observed in the Arabidopsis *upl3* mutant siliques.

*GL2* also negatively regulates seed lipid content by suppression of *PHOSPHOLIPASE D ALPHA 1* (*PLDα1*) gene expression (Liu et al., 2014). However, no significant difference in the expression of *PLDɑ1* was observed in WT and *upl3* mutant seeds (data not shown). Therefore, UPL3 may target other proteins for degradation during seed maturation that influence seed size and storage reserve accumulation.

### *upl3* mutants display increased expression of seed maturation genes

Several genes influence both seed lipid content and seed size in Arabidopsis, including *APETALA 2 (AP2)(Ohto et al., 2009), LEAFY COTYLEDON 1 (LEC1)* and *LEAFY COTYLEDON 2 (LEC2)* (Santos Mendoza et al., 2005). The expression of these genes was assessed during the development of *upl3* and WT seeds using q-RT-PCR. No differences in the expression of *AP2* (Figure 3A) or *LEC2* (Figure 3B) were seen between *upl3* and WT seeds. However, significant increases in *LEC1* expression were observed in *upl3* mutant seeds at 10 DPA compared to WT (Figure 3C). Transcription of *LEC1* is positively regulated by *LEC2* (Santos-Mendoza et al., 2008b; Baud et al., 2016). Given the observed increase in *LEC1*, but not *LEC2*, expression, it is possible that altered *UPL3* expression may affect LEC2 protein levels, thus altering *LEC1* expression in *upl3* mutant seeds. This was tested by measuring expression of *WRINKLED 1* (*WRI1*) and *MYB118,* which are also regulated by LEC2 (Barthole et al., 2014; Baud et al., 2009). Increased expression of both genes was observed in *upl3* mutant siliques relative to WT from 10 DPA (Figures 3D and 3E), supporting the hypothesis that UPL3 may influence LEC2 protein levels and expression of target genes.

### *UPL3* reduces LEC2-mediated transcription of seed maturation genes

LEC1 and LEC2 bind to the promoters and activate the expression of seed maturation genes, such as *OLEOSIN 1* (*OLE1*), a gene required for seed lipid accumulation (Santos-Mendoza et al., 2008b; Baud et al., 2016). To further assess the potential role of *AtUPL3* in LEC2*-*mediated gene expression, expression of *OLE1* in maturing seeds was measured in WT and *upl3* mutant Arabidopsis. Figure 3F showed that *OLE1* was expressed at higher levels in the *upl3* mutant, consistent with the hypothesis that UPL3 may influence LEC2 protein levels. To assess if UPL3 directly affects LEC2-mediated transcription of *OLE1*, transient expression of the Arabidopsis *OLE1* promoter fused to firefly luciferase was carried out in Arabidopsis *upl3* mesophyll protoplasts. The low levels of endogenous *OLE1* promoter activity (Figures 3G) were increased by co-transfection with *35S::LEC2*. Co-transfection with both *35S::LEC2* and *35S::UPL3* significantly reduced LEC2-induced *OLE1* promoter activity, suggesting that UPL3 reduces LEC2-mediated transcriptional regulation of seed lipid biosynthetic genes.

### UPL3 mediates the proteasomal degradation of LEC2

HECT E3 ligases such as UPL3 mediate the proteasomal degradation of substrate proteins by ubiquitylation (Maspero et al., 2013). To establish if UPL3 mediates the proteasomal degradation of LEC2, the stability of bacterially-expressed and purified LEC2-HIS was assessed in crude protein extracts of 10 DPA siliques of WT and *upl3* mutant Arabidopsis. Figure 4A shows that LEC2-HIS was unstable in WT silique protein extracts, and this instability was reduced by the proteasomal inhibitor MG132. In contrast, LEC2-HIS levels were more stable in protein extracts from *upl3* mutant siliques, and MG132 further reduced this instability. To support these observations of LEC2-HIS stability, LEC2 protein levels in Arabidopsis *upl3* mutant plants, in WT plants, and in transgenic plants expressing *35S::UPL3-HA* were assessed using LEC2-specific antibodies. Increased endogenous LEC2 protein levels were observed in mutant *upl3* siliques compared to WT plants (Figure 4B). Consistent with these observations, reduced LEC2 levels were seen in siliques of 35S*::3HA-UPL3* transgenic lines compared to WT siliques (Figure 4C). This indicated that *AtUPL3* influences LEC2 protein levels *in vivo*, and that LEC2-HIS has reduced proteolytic instability in the absence of *AtUPL3*.

### *UPL3* physically interacts with *LEC2* and promotes its ubiquitylation

Ubiquitylation by HECT E3 ligases involves the direct interaction of the E3 ligase with substrate proteins (Yau and Rape, 2016). A transient expression system was used to co-express 35S::*LEC2-GFP* and *35S::AtUPL3-FLAG* in *Nicotiana benthamiana* leaves. LEC2-GFP was co-purified with 3FLAG-UPL3 purified on FLAG beads (Figure 4D), demonstrating an interaction between 3FLAG-UPL3 and LEC2-GFP. Measurement of endogenous LEC2 protein in developing siliques taken from 35S::*UPL3* transgenic plants revealed LEC2 forms with higher apparent MW in comparison to those observed in WT protein extracts (Figure 4E). In transfected *Nicotiana benthamiana* leaves, increased levels of apparent higher MW LEC2-GFP forms were observed when 3FLAG-UPL3 was co-expressed with LEC2-GFP (Figure 4F). Western-blots of the same samples with anti-ubiquitin antibodies identified the apparent higher MW forms of LEC2-GFP as ubiquitylated LEC2-GFP. Taken together, these results provide evidence that UPL3 physically interacts with and promotes ubiquitylation of LEC2.

### Variation in the promoter of *BnaUPL3.C03* is sufficient for differential expression and influences variation in yield traits

We observed major differences in *BnaUPL3.C03* expression between GWAS accessions displaying variation in seed yield (Figure 2B and Supplementary Table 2). Alignment of *BnaUPL3.C03* promoter sequences (from 2 kb upstream of the ATG initiation codon) from a high-(Coriander) and a low-expressing (Dimension) accession revealed multiple sequence differences, including 34 Single Nucleotide Polymorphisms (SNPs) and 7 indels between 3-60nt (Supplementary Figure 6). The Dimension and Coriander *BnaUPL3.C03* promoters were fused to a firefly luciferase reporter gene (Figure 5A) and their activities assessed following protoplast transfection. Figure 5B shows that the Coriander promoter drove three times more luciferase activity than the Dimension promoter in Arabidopsis leaf protoplasts. These three-fold differences in *BnaUPL3.C03* expression were consistent with RNAseq and q-RT-PCR data (Figures 1E and 1F) from *B. napus* seedlings, showing that variation in promoter activity is the primary source of variation in *BnaUPL3.C03* expression between these two accessions.

**Figure 6.**
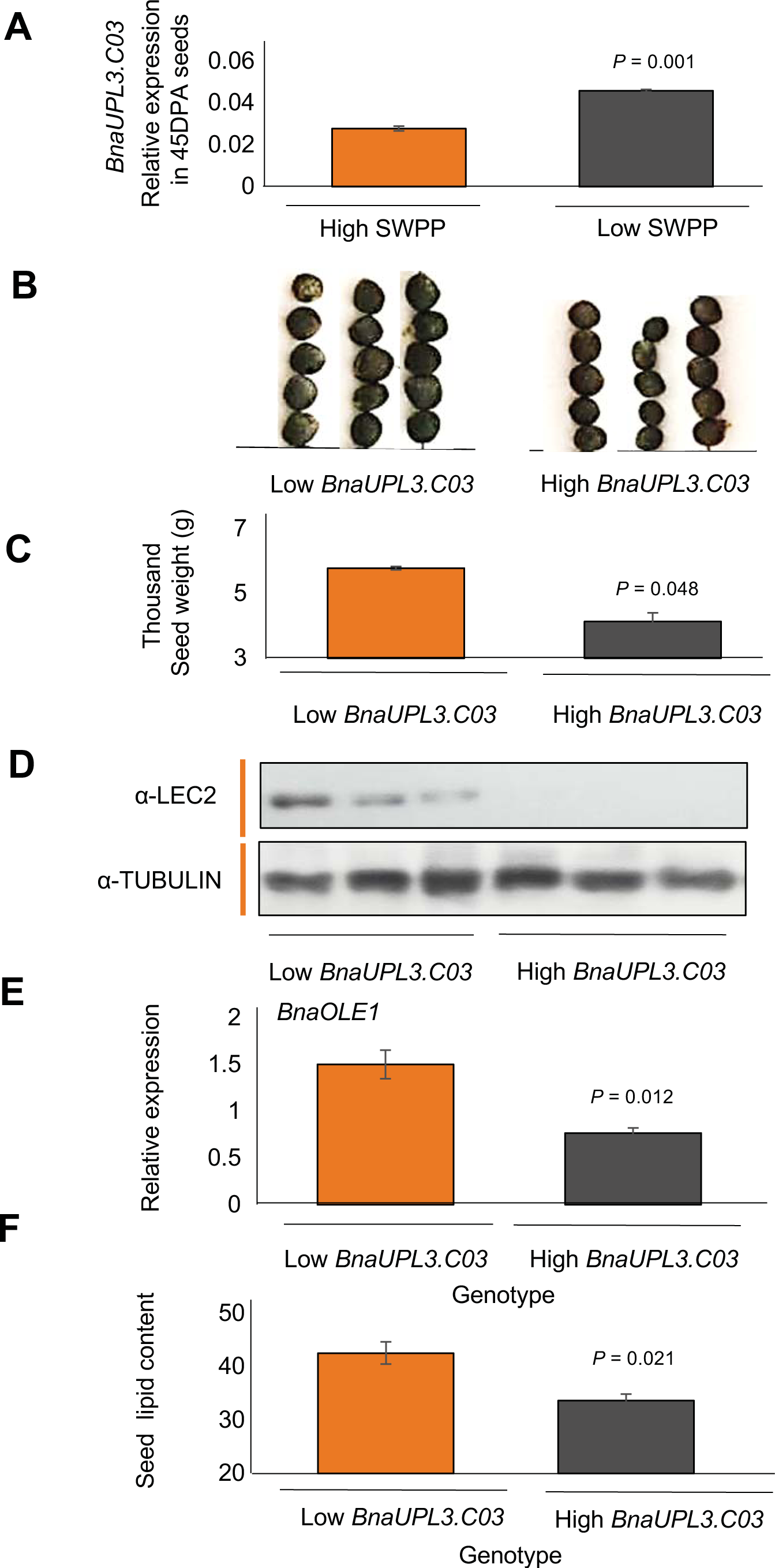
Relationships between *BnaUPL3.C03* expression levels in high- and low-SWPP *B. napus* accessions to seed size, seed LEC2 protein levels, and seed lipid content. **A.** *BnaUPL3.C03* expression levels in 45 DPA seeds in *B. napus* Dimension (DIM) with high SWPP, and Coriander accessions with low SWPP measured by q-RT-PCR. Expression levels are relative to the *BnaACTIN2* gene. Data are presented as means ± SE where n=3 for each genotype. *P* values were determined by Student’s *t*-test. **B.** Seed sizes in the low-expressing *BnaUPL3.C03* line Dimension and the high expressing *BnaUPL3.C03* line Coriander. **C.** Thousand seed weights of low-expressing *BnaUPL3.C03* line Dimension and the high-expressing *BnaUPL3.C03* line Coriander. Data are shown as means ± SE. Seeds were weighed in batches of 100 seed and thousand seed weight calculated based on these values. n=3 for each genotype included. *P* values were determined by Student’s *t*-test. **D.** LEC2 protein levels in 45 DPA seeds of three low *BnaUPL3.C03* expressing, and three high *BnaUPL3.C03* expressing accessions (Supplementary Table 2). Western blots of seed protein extract were probed with anti-LEC2 (top panel) and with anti-tubulin (lower panel) as a protein loading control. **E.** Expression of *BnaOLE1* was quantified by q-RT-PCR in low expressing *BnaUPL3.C03* (Dimension) and high expressing *BnaUPL3.C03* (Coriander) accessions. Expression levels are relative to the *BnaACTIN2* gene. n=3 for each genotype. Primers were designed to measure expression of both *BnaC01g17050D* and *BnaA01g14480D BnaOLE1* genes. *P* values were determined by Student’s *t*-test. **F.** Lipid content of mature seeds of low expressing *BnaUPL3.C03* (Dimension) and high expressing *BnaUPL3.C03* (Coriander) accessions. *P* values were determined by Student’s *t*-test.

To assess the role of *BnaUPL3.C03* promoter variation in variation in seed size, transgenic Arabidopsis lines expressing the Arabidopsis *UPL3* coding region driven by Coriander and Dimension *BnaUPL3.C03* promoters were constructed. Transgenic lines were made in the *upl3* mutant background to assess differential complementation of *upl3* mutant phenotypes in response to transgene expression. *UPL3* expression was measured in 10 DPA siliques of transgenic lines, the *upl3* mutant, and WT plants. The two *B. napus* promoters expressed the *AtUPL3* coding regions at the predicted levels in developing siliques (Figure 5C). Seed sizes in the high-expressing Coriander promoter transgenic lines showed near wild-type seed sizes, indicating complementation of the large Arabidopsis seed *upl3* phenotype. Negligible complementation of the large seed *upl3* phenotype was observed in plants expressing *AtUPL3* under the control of the low-expressing Dimension promoter (Figure 5D). Thus, variation in *BnaUPL3.C03* promoter activity can influence final seed size by altering levels of *AtUPL3* coding region expression.

### Differential expression of *BnaUPL3.C03* leads to variation in BnaLEC2 protein levels and determines final seed lipid content

Associative transcriptomics identified a negative relationship between *BnaUPL3.C03* expression and SWPP in *B. napus* accessions (Figure 6A). Consistent with seed size phenotypes seen in transgenic Arabidopsis (Figure 5D), low *BnaUPL3.C03* expressing lines had larger seeds (Figure 6b) and higher thousand seed weight (Figure 6c). To relate these *B. napus* phenotypes to the proposed mechanism of UPL3-mediated ubiquitylation and destabilization of LEC2 in Arabidopsis (Figure 4), LEC2 antibody was used to screen a subset of six *B. napus* accessions varying in *BnaUPL3.C03* expression (Supplementary Table 2 and Figure 2B). Western blots of protein extracted from 45 DPA seeds of these accessions were probed with anti-LEC2 antibodies to assess variation in LEC2 protein levels. Figure 6D shows that LEC2 protein levels were higher in all three *B. napus* accessions with reduced *BnaUPL3.C03* expression, compared to accessions with higher *BnaUPL3.C03* expression, in which no LEC2 was detected in seeds using this method. These observations show that the mechanism of *UPL3*-mediated control of LEC2 protein levels established in Arabidopsis underlies variation in seed size observed in *B. napus* accessions. As predicted by the mechanism, significant differential expression of *BnaOLE1* was observed across the subset of *B. napus* accessions displaying differential expression of *BnaUPL3.C03* during seed maturation (Figure 6E). Finally, accessions displaying reduced *BnaUPL3.C03* and consequent increased *BnaOLE1* expression levels exhibit significantly higher seed lipid levels relative to those displaying high *BnaUPL3.C03* expression (Figure 6F).

### Reduced *BnUPL3.C03* expression has not yet been exploited in current *B. napus* breeding material

To assess if variation in *BnaUPL3.C03* expression levels has been selected for in the breeding of current elite oilseed rape varieties, expression was measured across a panel of seven current elite *B. napus* lines. Expression levels were compared to those measured in GWAS accessions exhibiting high differential expression of *BnaUPL3.C03* and high variation in yield traits (Figure 7). There was substantial variation in *BnaUPL3.C03* expression in the elite lines, with relatively high levels of expression in several lines. Therefore, there is significant potential for further yield increases in elite oilseed rape germplasm through reduction of *BnaUPL3.C03* expression.

**Figure 7.**
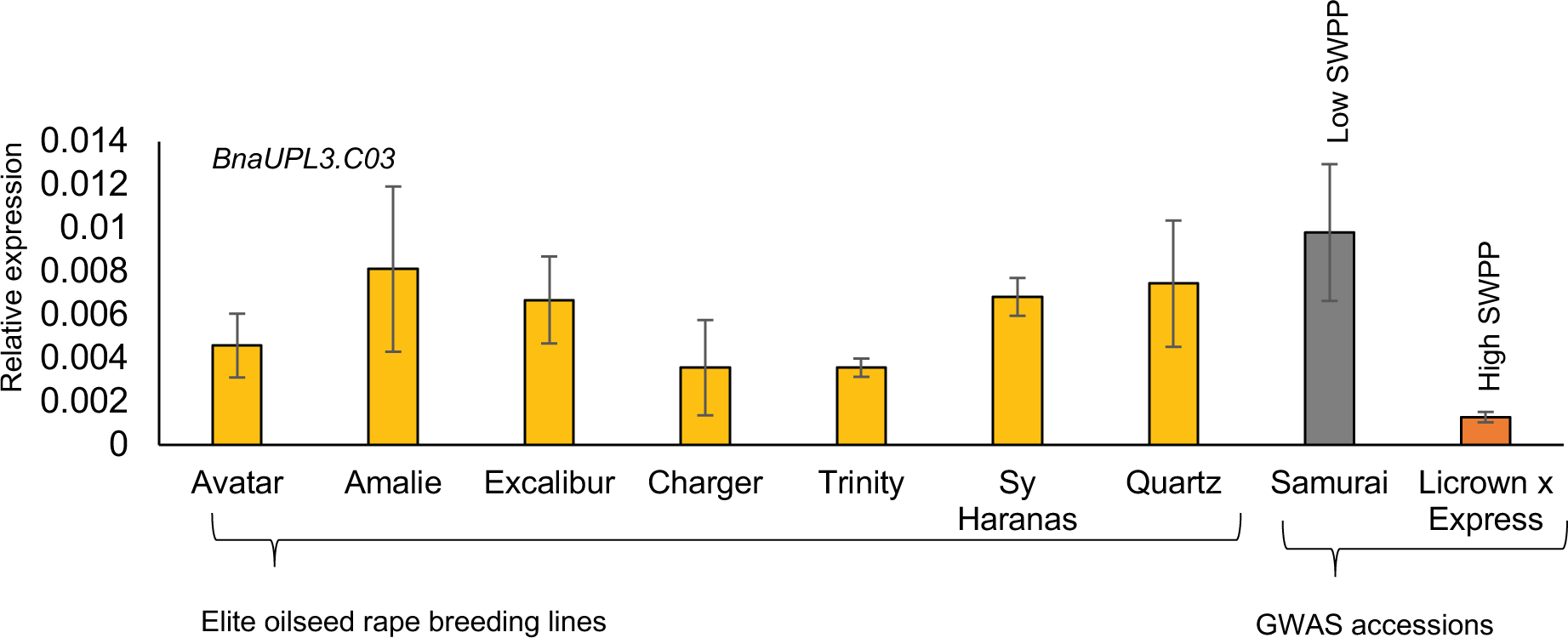
Selection for low *BnaUPL3.C03* expression has not yet been exploited in elite breeding lines. *BnaUPL3.C03* expression levels in 45 DPA seeds of seven current elite commercial cultivars of oilseed rape were measured using q-RT-PCR and compared with expression levels of *BnaUPL3.C03* from GWAS accessions Samurai and Licrown x Express, which exhibit high and low *BnaUPL3.C03* expression levels, and low and high SWPP phenotypes, respectively. *BnaACTIN2* gene expression was used for comparison using q-RT-PCR control. Data are presented as means ±SE (n=3 for each genotype).

## Discussion

Using Associative Transcriptomics (Harper et al., 2012) we identified variation in the expression of a *B. napus* gene, *BnaUPL3.C03*, that modulates seed size, lipid content and field yields in this important oilseed crop. *BnaUPL3.C03* encodes an orthologue of the Arabidopsis HECT E3 ligase *UPL3* gene. We show that it promotes the ubiquitylation and destabilization of LEC2, a “hub” transcriptional regulator of seed storage processes (Boulard et al., 2017). Reduced *BnaUPL3.C03* expression maintains higher levels of LEC2 during seed maturation, prolonging transcriptional activation of storage lipid genes, leading to larger seeds with elevated lipid levels in *B. napus.* Lines with relatively low *BnaUPL3.C03* expression had robust yield increases in field conditions compared to lines with higher *BnaUPL3.C03* expression. Variation in *UPL3* expression levels in elite oilseed rape cultivars identifies a promising approach for achieving further increases in oilseed yields by selecting for reduced *UPL3* expression levels.

Comparison of assemblies of the *BnaUPL3.C03* gene and flanking sequences from two *B. napus* lines with low and high *BnaUPL3.C03* expression identified high levels of promoter sequence variation that segregated with high SWPP and reduced *BnaUPL3.C03* expression (Figure 5B and 5C). Transient assays showed that a 2kb 5’ flanking region from high- and low-expressing *BnaUPL3.C03* alleles promoted the expected expression differences (Figure 5B). Driving expression of the Arabidopsis *UPL3* coding sequence with *B. napus* promoter variants led to differential complementation of the Arabidopsis *upl3* mutant seed phenotypes (Figure 5D). This established that natural variation in the activity of the *BnaUPL3.C03* promoter is sufficient to cause variation in seed size and yield traits. The multiple variants detected between two promoters driving differential expression and the continuous profile of *BnaUPL3.C03* expression in the accessions (Figure 1E) suggests there may be multiple sequence variants that together reduce promoter activity. Finally, variation in other regions of the *BnaUPL3.C03* gene, including the coding region, may also have the potential to contribute to variation in transcript abundance and yield observed in *B. napus* varieties.

Such genetic variation influencing gene regulation is an increasingly important resource for trait improvement, including increasing seed yields. For example, natural variation in the copy number of a promoter silencing element of the FZP gene underlies variation in spikelet numbers in rice panicles and is an important determinant of yield (Bai et al., 2017). Similarly, QTL analyses identified promoter variation in the rice *GW7* gene that, combined with variation that reduced expression of an *SPL16* transcriptional repressor of *GW7,* led to 10% increases in grain yield and improved quality(Wang et al., 2015). A deletion in the promoter region of *GW5* in Nipponbare rice lines reduced expression and increased grain width (Liu et al., 2017), and similarly, promoter deletions in *GSE5* accounted for wide grains in *indica* rice varieties (Duan et al., 2017). More generally, genetic variation in chromatin accessibility (a mark of promoter activity) explained about 40% of heritable variation in many quantitative traits in maize (Rodgers-Melnick et al., 2016). These reports, and the study described here, reveal the exceptional promise of accessing variation in promoter sequences and altered transcriptional activity for identifying new regulatory mechanisms and for the quantitative manipulation of complex traits such as yield in crop plants.

UPL3 was first identified in Arabidopsis as a loss-of-function mutation in a HECT E3 ligase gene causing increased leaf hair branching (Downes et al., 2003; El Refy et al., 2003). *UPL3* expression increased during seed maturation in Arabidopsis (Figure 2A), and an ortholog, *BnaUPL3.C03,* was differentially expressed in the pods of *B. napus* lines (Figure 2B) that varied in seed weight per pod (Figure 1A). LEC2, a transcriptional regulator of seed storage processes, is more stable in a *upl3* loss-of-function mutation in Arabidopsis (Figure 4A) and UPL3 promoted the ubiquitylation and destablization of LEC2 (Figures 3E and 3F). Consistent with these observations, in maturing seeds of *B. napus* lines with low *BnaUPL3.C03* expression, LEC2 protein levels remained higher than in lines displaying higher *BnaUPL3.C03* transcript levels (Figure 6D). These observations suggested that UPL3 ubiquitin ligase activity modulates levels of LEC2 protein during seed maturation. *LEC1* is directly activated by LEC2, and together they activate the expression of genes involved in promoting seed lipid accumulation, such *OLE1.* Consistent with this, we observed elevated *OLE1* expression levels in *upl3* mutant seeds relative to WT and showed that *BnaOLE1* is differentially expressed between *B. napus* accessions displaying differential expression of *BnaUPL3.C03*. These observations show that reduced levels of UPL3 maintain higher levels of LEC2 protein during seed maturation, thus prolonging the duration of expression of seed maturation genes, leading to increased seed lipid in Arabidopsis *upl3* mutants and *B. napus* accessions with reduced *BnaUPL3.C03* expression.

At earlier stages of seed development, the seed coat in Arabidopsis *upl3* mutants has altered mucilage and reduced expression of *GL2* (Figure 2F), a transcription factor controlling epidermal cell differentiation(Lin et al., 2015). GL2 promotes expression of the rhamnose biosynthesis gene *MUM4* that is required for seed mucilage production(Lin et al., 2015; Shi et al., 2012), and expression of *MUM4* is also reduced during testa development in *upl3* (Figure 2G). This observation revealed a unifying role of *UPL3* in regulating both testa and embryo maturation by modulating levels of transcription factors during different stages of seed development. These transcription factors, GL3 and EGL3 (Patra et al., 2013), and LEC2 (this study) in turn modulate expression of other transcription factors and biosynthetic genes involved in testa and embryo development.

Altered expression of *LAFL* genes has profound developmental consequences such as ectopic embryogenesis (Stone et al., 2001; Roscoe et al., 2015), but induced expression of *LEC2* in vegetative tissues does increase lipid accumulation (Andrianov et al., 2010; Santos Mendoza et al., 2005). These studies show that the activities of *LEC2* expression in storage processes and embryo development are difficult to separate, probably due to the timings of expression, interdependence and partial redundancy of *LAFL* gene function. By identifying a mechanism controlling LEC2 protein levels during seed maturation, we have shown that it is possible to elevate lipid levels during normal embryo development (Figure 2D). Mis-expression of *WRI1* permits normal seed development while increasing lipid content (Kanai et al., 2016; van Erp et al., 2014) by extending seed maturation, consistent with our observations of elevated *WRI1* expression in *upl3* (Figure 3E). Intensive breeding is optimizing oilseed lipid composition for different end-uses, but comparatively slow progress is being made in increasing yields of oilseed crops such as oilseed rape, with current rates of yield increase predicted to be insufficient to meet future needs (Ray et al., 2013). Variation in the promoter of *BnaUPL3.C03* that reduces expression and increases seed lipid content has not yet been exploited in oilseed rape breeding (Figure 7), demonstrating how lipid content and seed yields could be increased without influencing composition. The presence of a potential LEC2 ortholog in soybean (Manan et al., 2017), and the expression of *LEC2* and other B3-domain transcription factor homologs during seed lipid synthesis in sunflower (Badouin et al., 2017) and oil palm (Singh et al., 2013) reveals the potential of UPL3-mediated regulation of LEC2 to increase seed lipid levels and overall yields in other major oilseed crops.

## Methods

### Plant material and growth conditions

Phenotype data was collected from 94 accessions representing winter, spring and Chinese oilseed rape from the OREGIN *Brassica napus* fixed foundation diversity set(Harper et al., 2012). Plants were grown in a randomised, triplicated experimental design in a Kedar greenhouse under natural light with no controlled heating. Prior to transplantation, plants were grown (18/15°C day/night, 16hr light) for 4 weeks before 6 weeks vernalisation (4°C, 8hr light). 20 typical pods were collected from each mature plant and digitally imaged. Pod length (Podl) was measured using ImageJ (Schneider et al., 2012). Pods were weighed (PW) before threshing to remove seeds. Seed numbers, average seed length (SL), width (SW), area (SA), single seed weight (SSW) and thousand grain weight (TGW) were measured for each sample using a Marvin device (GTA Sensorik GmbH, Germany). Numbers of seeds per pod (SPP), seed weight per pod (SWPP) and seed density (SDen) were calculated from these data. Field yields of selected accessions were grown in four replicated field plots (1.25m x 6m, Church Farm, Bawburgh, Norfolk, UK) in a randomized design and harvested by combine. Total plot yield was determined for each plot and an average plot yield taken.

All *Arabidopsis thaliana* mutant and transgenic lines used were in Col-0 background. Plants were grown on soil in a growth chamber with 16/8 hr day/night at 22°C after 48 hours stratification at 5°C. A sequence indexed T-DNA insertion line in At4G38600 (UPL3), Salk_015334, was obtained from The Nottingham Arabidopsis Stock Centre (NASC). Genotyping primers were designed using the primer design tool http://signal.salk.edu/tdnaprimers.2.html: Primer sequences are in Supplementary Table 2. Genotyping used TAKARA EX taq (Takara Bio, USA).

To identify loss-of-function mutations in *UPL3* in *Brassicas*, a TILLING population (Stephenson et al., 2010) of *B. rapa* (the A genome donor to *B. napus*) was screened for mutations in the predicted gene Bra010737 on chromosome A08 that encodes the single ortholog of *Arabidopsis thaliana UPL3*. The predicted *B. rapa* gene was defined by a full-length transcript Bra010737.1. There was only one clear ortholog of *UPL3* in *B.rapa*, consistent with two copies present in the amphidiploid *B. napus* on chromosomes A08 and C03. Two premature stop codon mutations were identified in Bra010737 (Supplementary Figure 7) in lines JI32517 and JI30043. Primers were designed to amplify genomic DNA from the mutated region (Supplementary Table 4) and used to validate the TILLING mutations. Line JI32517-B was validated and used in further studies. The line was selfed and progeny screened for homozygous (G>A) and “WT” (C>T) changes by sequencing of the locus. Seeds were harvested from the “WT” and *upl3* mutant and their area measured using ImageJ.

A subset of *B. napus* accessions and a panel of elite *B. napus* breeding lines were grown under glasshouse conditions after vernalisation. Leaf material was harvested from the first true leaf and stored at −70℃ prior to further processing. Developing pods were staged by tagging when petals were beginning to emerge from the developing bud, taken as zero days post anthesis (DPA). Samples for RNA isolation were collected at 45 DPA and stored at −70℃. For expression analyses in Arabidopsis, WT and *upl3* mutant plants were grown as described, without vernalization, and floral buds tagged when petals were beginning to emerge from the developing bud (0 DPA). Siliques were then harvested at 0, 5, 10 and 15 DPA and tissue samples stored at −70℃.

### Population structure analysis

Associative Transcriptomics analysis was carried out as described(Harper et al., 2012). The population structure Q matrix was re-calculated using unlinked markers across the set of 94 lines. One SNP per 500 kb interval along pseudomolecules, excluding regions less than 1000 kb from centromeres (Mason et al., 2017), were selected for Bayesian population structure analysis via STRUCTURE 2.3.3 (Pritchard et al., 2000). This analysis incorporated a minor allele frequency of 5%. The optimum number of K populations was selected as described (Harper et al., 2012).

### SNP analysis

SNP data, STRUCTURE Q matrix and phenotypes of the 94 accessions were combined using TASSEL (V4.0). Following the removal of minor alleles (frequency <0.05) ~144,000 SNPs were used to calculate a kinship (K) matrix to estimate the pairwise relatedness between individuals. Data sets were analysed using both Generalised and Mixed Linear Models (GLM and MLM). Goodness of fit of the model was determined by a QQ plot of the observed versus the expected −log10P values. −log10P values were plotted in chromosome order and visualized using R programming scripts (http://www.R-project.org/.)

### GEM analysis

Relationships between GEM sequence read depth (RPKM) and trait data variation were analysed by Linear regression analysis using R. −log10P values were plotted in pseudomolecule order and visualized using R scripts.

### DNA constructs

The *p35S::3HA-AtUPL3* transgenic line was generated by cloning *AtUPL3* cDNA into the pENTR TOPO-D vector (Thermofisher, UK) using the primers described in Supplementary Table 4. LR Clonase mix II (Thermofisher, UK) was used to transfer the *AtUPL3* CDS into the 35S PB7HA binary vector. All constructs were sequenced before use. The *p35S::3HA-AtUPL3* construct was transformed into *Agrobacterium tumefaciens* strain GV3101, and Arabidopsis *Col-0* plants were transformed using the floral dip method (Clough and Bent, 1998).

Promoters of the *BnaUPL3.C03* gene from the high- and low-expressing accessions Coriander and Dimension were amplified using primers described in Supplementary Table 4. *AtUPL3* CDS in the pENTR TOPO-D vector was transferred into the binary vector pEarley201 using LR Clonase, and *BnaUPL3.C03* promoter PCR products, digested with Stu1 and Xho1, were ligated into the pEarly 201-*AtUPL3* CDS plasmid. The *BnaUPL3.C03* promoter::*AtUPL3* CDS construct was transformed into *Agrobacterium tumefaciens* strain GV3101, and Arabidopsis *upl3* mutant plants were transformed using the floral dip method (Clough and Bent, 1998).

### Arabidopsis and *Brassica* seed size quantification

Arabidopsis seeds were harvested from mature plants and the seeds imaged at 10x magnification. *B. napus* and *B.rapa* seeds were harvested from mature plants and scanned using a photocopier. Seed area was quantified using ImageJ particle analysis.

### Ruthenium red staining

Seed mucilage phenotypes were assessed using methods described by (McFarlane et al., 2014). Stained seeds were imaged at 10x magnification.

### Seed lipid quantification and profiling

Fatty acid profile analyses in Arabidopsis were carried out using the methods described by (Li et al., 2006). Lipid content of *B. napus* seeds were measured using Near Infrared Spectroscopy (Wang et al., 2014).

### PCR and sequencing

All PCR reactions were carried out using Phusion High Fidelity DNA polymerase (New England BioLabs) according to manufacturer's instructions. Capillary sequencing was carried out by GATC Biotech (Germany).

### cDNA synthesis and q-RT-PCR

RNA was extracted using the SPECTRUM Total Plant RNA kit (Sigma, UK). 1ug of RNA was treated with RQ1 RNase-Free DNase (Promega, USA) and cDNA synthesis was carried out using the GoScript Reverse Transcription system (Promega, USA) using OligoDT. All protocols were carried out using manufacturers’ guidelines. cDNA samples were diluted 1:10 in water before use. Q-RT-PCR was carried out using SYBR green real time PCR mastermix (Thermofisher) and performed using Lightcycler 480 (Roche, Switzerland). Primer sequences used for q-RT-PCR are in Supplementary Table 4. Primer efficiencies and relative expression calculations were performed according to methods described (Pfaffl, 2001). All q-RT-PCR assays were repeated at least twice.

### Promoter transactivation assay

Promoters and full length cDNAs of selected Arabidopsis Col-0 genes were amplified by PCR using Phusion polymerase (Thermo Scientific) according to the manufacturer's guidelines. Promoter primer sequences are in Supplementary Table 2. PCR reactions were purified using Wizard SV Gel and PCR Clean up system (Promega, UK) and inserted into pENTR D-TOPO vector (Thermofisher, UK), and an LR reaction was used to clone promoters into a Firefly Luciferase reporter vector fLUC (pUGW35). *LEC2* and *UPL3* CDS were transferred using LR Clonase into PB7HA and PW1266 respectively to create *p35S::3HA-LEC2* and *p35S::3FLAG-UPL3*. A *35S::Renilla* luciferase construct was used to quantify relative promoter activities. Plasmid preparations for transient assays were prepared using the Qiagen Plasmid Maxi Kit according to manufacturer’s instructions.

Promoter transactivation assays were carried out using protoplasts isolated from *upl3* leaves (Wu et al., 2009). Assays were carried in triplicate using 5ug of plasmid and 100ul of purified protoplasts (approximately 50,000 cells). After an overnight incubation at room temperature, transfected protoplasts were harvested and promoter activity assessed using the Dual Luciferase Reporter assay system (Promega, USA). The ratio of Firefly Luciferase to Renilla Luciferase was determined using the dual assay Promega protocol on a Glomax 20/20 luminometer (Promega, USA). All transactivation assays were repeated at least twice.

### Total protein extraction from *B. napus* pods and Arabidopsis siliques

Material was ground to a fine powder in liquid nitrogen and resuspended in extraction buffer (1ml/g fresh weight) (25mM Tris-HCl pH 8.0, 10mM NaCl, 10mM MgCl_2_, 4mM AEBSF and 50µM MG132). Following an incubation on ice for 30 minutes, samples were centrifuged at maximum speed for 5 minutes at 4℃. Supernatant was then added to a fresh tube and centrifugation repeated for 10 minutes. Total protein content was assessed using Bradford reagent (Bio-Lab) according to manufacturer's instructions.

### Protein expression in *E. coli* BL21

Arabidopsis LEC2 CDS was amplified from cDNA using primers described in Supplementary Table 4. Following purification, PCR products were digested with and ligated into digested pET-24a vector. Plasmids were transformed into BL21 *E. coli* cells. Following overnight growth, a single colony was used to inoculate 10ml liquid LB+ kanamycin 25 μg/ml and incubated at 37°C overnight. The following day, 4ml of this culture was used to inoculate 400ml LB+kanamycin and incubated for 2 hours at 37°C. IPTG was then added to 100μM and the culture incubated at 28°C for 3 hours to induce protein expression. Cultures were then centrifuged at 3,500rpm for 10 minutes at 4°C and the cell pellet suspended in 7.5ml of suspension buffer (50mM HEPES pH 7.5, 150mM NaCl, 1% TritonX-100; 10% glycerol; 1 Roche EDTA-FREE inhibitor cocktail tablet). Cells were sonicated 4 x 10 seconds with 20 sec intervals on ice, and sonicates were centrifuged at 12,000g for 20 minutes at 4°C. Protein purification was carried out using Dynabeads ^®^ His-tag magnetic beads (Novex). Prior to use, beads were washed 3 times in wash buffer (50 mM HEPES (PH7.5); 150 mM NaCl; 10% glycerol). The sonicate was then added to washed beads and incubated at 4°C with rotation for 30 minutes. Beads were then washed 3 times with suspension buffer and 3 times with wash buffer. His-tagged protein was eluted from beads using 120μl elution buffer (300mM Imidazole, pH 7.5, 1x PBS pH 7.5, 0.3M NaCl, 0.1% Tween-20, 10% glycerol). Purified protein was quantified using Bradford reagent (Bio-Rad) and stored at −70°C in 15μl aliquots.

### Cell-free degradation assay and western blot analyses

Total protein was extracted from 10-15 DPA siliques taken from WT and *upl3* mutant plants using the following extraction buffer: 25mM Tris-HCL pH7.5, 10mM NaCl, 10mM MgCl_2_ and 4mM AEBSF. Following protein quantification using BioRad reagent (Bio-Lab), 20µg total protein was added to degradation buffer (25mM Tris-HCL, 10mM MgCl_2_, 5mM DTT, 10mM NaCl, 10mM ATP and 5µg *E.coli*-expressed LEC2-HIS). 30µl samples were incubated at 22℃ for 0, 5, 10 and 15 minutes in the presence or absence of 200 μM MG132. Reactions were stopped with 4x SDS Laemmli sample buffer (Bio-Rad) and samples denatured at 96°C for 5 minutes. Samples were loaded on 4-20% gradient SDS gels and probed with anti-HIS antibody to visualise LEC2-HIS protein. LEC2 protein was also measured using purified rabbit polyclonal antibodies raised against the antigenic peptide ARKDFYRFSSFDNKKL from LEC2 coupled to keyhole limpet haemocyanin (New England Peptides, USA). This assay was repeated three times. Western blots used antibodies at the following dilutions: anti-FLAG 1/1000; anti-GFP 1/5000; anti-tubulin 1/5000; anti-HA 1/1000; anti LEC2 1/1000. Secondary antibodies used for LEC2 were anti-rabbit 1/5000; for tubulin anti-mouse 1/5000. Western blots were developed with FEMTO Max peroxidase substrate (Fisher Scientific).

### Transient expression in *Nicotiana benthamiana*

The Arabidopsis *UPL3* cDNA TOPO construct cloned into the 3xFLAG PW1266 vector. Full length Arabidopsis LEC2 cDNA was amplified using primers described in Supplementary Table 4. Following cloning into pENTR TOPO-D, *LEC2* CDS was transferred by LR reaction to pEARLEY 103. The resulting constructs, *p35S::3XFLAG-UPL3* and *p35S::LEC2-GFP* were transformed into *Agrobacterium tumefaciens* GV3101 and 10 ml cultures grown at 28°C for 48 hours before transfection into leaves of 4 week old *Nicotiana benthamiana* plants using a 1 ml syringe without a needle. *Agrobacterium* cultures were transfected into the underside of the leaves and after 48 hours, transfected regions were excised and stored in liquid nitrogen before purification. Protein extracts were purified using M2 FLAG beads (Sigma). Purified protein was eluted from beads using 1X SDS buffer and heated at 96C for 5 minutes. Samples were loaded onto a 4-20% gradient SDS gel and western blotted with anti-GFP antibody or anti FLAG antibodies to assess 3FLAG-UPL3-LEC2-HIS interactions. For assessing LEC-GFP ubiquitylation in transient assays, total protein was extracted after transfection for 48 hrs and Western blotted using anti-GFP and then anti-ubiquitin. To detect in vivo ubiquitylation of LEC2, total protein was extracted from pooled 10-15 DPA siliques of a 35S::*AtUPL3-HA* transgenic line and from WT plants. Transgene expression was assessed following purification on Pierce Anti-HA magnetic beads (Thermo Scientific) and total protein extracts probed using anti-LEC2 to assess LEC2 ubiquitylation. Protein samples in SDS sample buffer were electrophoresed on precast 12%, 20% or 4-20% gradient SDS polyacrylamide gels (RunBlue, Expedeon Ltd, Cambridge, UK), transferred to PVDF membranes (Roche Diagnostics, Burgess Hill, UK) and immunoblotted. PVDF membranes were washed for 10 min in 50 ml PBS after transfer, then treated with blocking solution (5% w/v milk powder, 0.1% v/v Tween-20 in PBS) for an hour. Primary antibodies were diluted in blocking solution and incubated with the membrane for 1 h before 5 washes with PBST (PBS with 0.1% v/v Tween-20) at room temperature. Washed membranes were then treated with FEMTO Max peroxidase substrate (Fisher Scientific, Loughborough, UK) for 5 min before exposure to X-ray film.

## Acknowledgements

This work was supported by an ERA-CAPS grant (“ABCEED”) to MWB and LL. MWB was also supported by a Biological and Biotechnological Sciences Research Council (BBSRC) Institute Strategic Grants GRO (BB/J004588/1) and GEN (BB/P013511/1). LL is also supported by Labex Saclay Plant Sciences-SPS (ANR-10-LABX-0040-SPS). We thank Jingkun Ma for advice on transient expression and luciferase assays.

## Author Contributions

CM, MWB, RW, BD and LL designed the study; CM, NMcK, RW, JB and AF performed experiments, CM and MWB analyzed data and wrote the paper.

## References

Andrianov, V., Borisjuk, N., Pogrebnyak, N., Brinker, A., Dixon, J., Spitsin, S., Flynn, J., Matyszczuk, P., Andryszak, K., Laurelli, M., Golovkin, M., and Koprowski, H. (2010). Tobacco as a production platform for biofuel: overexpression of Arabidopsis DGAT and LEC2 genes increases accumulation and shifts the composition of lipids in green biomass. Plant Biotechnol. J. 8: 277–287.

Badouin, H. et al. (2017). The sunflower genome provides insights into oil metabolism, flowering and Asterid evolution. Nature 546: 148–152.

Bai, X., Huang, Y., Hu, Y., Liu, H., Zhang, B., Smaczniak, C., Hu, G., Han, Z., and Xing, Y. (2017). Duplication of an upstream silencer of FZP increases grain yield in rice. Nat Plants 3: 885–893.

Barthole, G., To, A., Marchive, C., Brunaud, V., Soubigou-Taconnat, L., Berger, N., Dubreucq, B., Lepiniec, L., and Baud, S. (2014). MYB118 represses endosperm maturation in seeds of Arabidopsis. Plant Cell 26: 3519–3537.

Baud, S. et al. (2016). Deciphering the Molecular Mechanisms Underpinning the Transcriptional Control of Gene Expression by Master Transcriptional Regulators in Arabidopsis Seed. Plant Physiol. 171: 1099–1112.

Baud, S., Wuillème, S., To, A., Rochat, C., and Lepiniec, L. (2009). Role of WRINKLED1 in the transcriptional regulation of glycolytic and fatty acid biosynthetic genes in Arabidopsis. Plant J. 60: 933–947.

Bevan, M.W., Uauy, C., Wulff, B.B.H., Zhou, J., Krasileva, K., and Clark, M.D. (2017). Genomic innovation for crop improvement. Nature 543: 346–354.

Boulard, C., Fatihi, A., Lepiniec, L., and Dubreucq, B. (2017). Regulation and evolution of the interaction of the seed B3 transcription factors with NF-Y subunits. Biochim. Biophys. Acta 1860: 1069–1078.

Chan, A., Carianopol, C., Tsai, A.Y.-L., Varathanajah, K., Chiu, R.S., and Gazzarrini, S. (2017). SnRK1 phosphorylation of FUSCA3 positively regulates embryogenesis, seed yield, and plant growth at high temperature in Arabidopsis. J. Exp. Bot. 68: 4219–4231.

Clough, S.J. and Bent, A.F. (1998). Floral dip: a simplified method for Agrobacterium-mediated transformation of Arabidopsis thaliana. Plant J. 16: 735–743.

Downes, B.P., Stupar, R.M., Gingerich, D.J., and Vierstra, R.D. (2003). The HECT ubiquitin-protein ligase (UPL) family in Arabidopsis: UPL3 has a specific role in trichome development. Plant J. 35: 729–742.

Duan, P., Xu, J., Zeng, D., Zhang, B., Geng, M., Zhang, G., Huang, K., Huang, L., Xu, R., Ge, S., Qian, Q., and Li, Y. (2017). Natural Variation in the Promoter of GSE5 Contributes to Grain Size Diversity in Rice. Mol. Plant 10: 685–694.

Duong, S., Vonapartis, E., Li, C.-Y., Patel, S., and Gazzarrini, S. (2017). The E3 ligase ABI3-INTERACTING PROTEIN2 negatively regulates FUSCA3 and plays a role in cotyledon development in Arabidopsis thaliana. J. Exp. Bot. 68: 1555–1567.

El Refy, A., Perazza, D., Zekraoui, L., Valay, J.-G., Bechtold, N., Brown, S., Hülskamp, M., Herzog, M., and Bonneville, J.-M. (2003). The Arabidopsis KAKTUS gene encodes a HECT protein and controls the number of endoreduplication cycles. Mol. Genet. Genomics 270: 403–414.

van Erp, H., Kelly, A.A., Menard, G., and Eastmond, P.J. (2014). Multigene engineering of triacylglycerol metabolism boosts seed oil content in Arabidopsis. Plant Physiol. 165: 30–36.

Fatihi, A., Boulard, C., Bouyer, D., Baud, S., Dubreucq, B., and Lepiniec, L. (2016). Deciphering and modifying LAFL transcriptional regulatory network in seed for improving yield and quality of storage compounds. Plant Sci. 250: 198–204.

Harper, A.L., Trick, M., Higgins, J., Fraser, F., Clissold, L., Wells, R., Hattori, C., Werner, P., and Bancroft, I. (2012). Associative transcriptomics of traits in the polyploid crop species Brassica napus. Nat. Biotechnol. 30: 798–802.

Kanai, M., Mano, S., Kondo, M., Hayashi, M., and Nishimura, M. (2016). Extension of oil biosynthesis during the mid-phase of seed development enhances oil content in Arabidopsis seeds. Plant Biotechnol. J. 14: 1241–1250.

Lin, Q., Ohashi, Y., Kato, M., Tsuge, T., Gu, H., Qu, L.-J., and Aoyama, T. (2015). GLABRA2 Directly Suppresses Basic Helix-Loop-Helix Transcription Factor Genes with Diverse Functions in Root Hair Development. Plant Cell 27: 2894–2906.

Liu, J. et al. (2017). GW5 acts in the brassinosteroid signalling pathway to regulate grain width and weight in rice. Nat Plants 3: 17043.

Liu, Y.-F. et al. (2014). Soybean GmMYB73 promotes lipid accumulation in transgenic plants. BMC Plant Biol. 14: 73.

Li, Y., Beisson, F., Pollard, M., and Ohlrogge, J. (2006). Oil content of Arabidopsis seeds: the influence of seed anatomy, light and plant-to-plant variation. Phytochemistry 67: 904–915.

Lu, K. et al. (2017). Genome-Wide Association and Transcriptome Analyses Reveal Candidate Genes Underlying Yield-determining Traits in Brassica napus. Front. Plant Sci. 8: 206.

Manan, S., Ahmad, M.Z., Zhang, G., Chen, B., Haq, B.U., Yang, J., and Zhao, J. (2017). Soybean LEC2 Regulates Subsets of Genes Involved in Controlling the Biosynthesis and Catabolism of Seed Storage Substances and Seed Development. Front. Plant Sci. 8: 1604.

Mason, A.S., Higgins, E.E., Snowdon, R.J., Batley, J., Stein, A., Werner, C., and Parkin, I.A.P. (2017). A user guide to the Brassica 60K Illumina Infinium^TM^ SNP genotyping array. Theor. Appl. Genet. 130: 621–633.

Maspero, E., Valentini, E., Mari, S., Cecatiello, V., Soffientini, P., Pasqualato, S., and Polo, S. (2013). Structure of a ubiquitin-loaded HECT ligase reveals the molecular basis for catalytic priming. Nat. Struct. Mol. Biol. 20: 696–701.

McFarlane, H., Gendre, D., and Western, T. (2014). Seed Coat Ruthenium Red Staining Assay. BIO-PROTOCOL 4.

Napier, J.A. and Graham, I.A. (2010). Tailoring plant lipid composition: designer oilseeds come of age. Curr. Opin. Plant Biol. 13: 330–337.

Nodine, M.D. and Bartel, D.P. (2010). MicroRNAs prevent precocious gene expression and enable pattern formation during plant embryogenesis. Genes Dev. 24: 2678–2692.

Ohto, M.-A., Floyd, S.K., Fischer, R.L., Goldberg, R.B., and Harada, J.J. (2009). Effects of APETALA2 on embryo, endosperm, and seed coat development determine seed size in Arabidopsis. Sex. Plant Reprod. 22: 277–289.

Patra, B., Pattanaik, S., and Yuan, L. (2013). Ubiquitin protein ligase 3 mediates the proteasomal degradation of GLABROUS 3 and ENHANCER OF GLABROUS 3, regulators of trichome development and flavonoid biosynthesis in Arabidopsis. Plant J. 74: 435–447.

Pfaffl, M.W. (2001). A new mathematical model for relative quantification in real-time RT–PCR. Nucleic Acids Res. 29: e45–e45.

Pritchard, J.K., Stephens, M., and Donnelly, P. (2000). Inference of population structure using multilocus genotype data. Genetics 155: 945–959.

Ray, D.K., Mueller, N.D., West, P.C., and Foley, J.A. (2013). Yield Trends Are Insufficient to Double Global Crop Production by 2050. PLoS One 8: e66428.

Rodgers-Melnick, E., Vera, D.L., Bass, H.W., and Buckler, E.S. (2016). Open chromatin reveals the functional maize genome. Proc. Natl. Acad. Sci. U. S. A. 113: E3177–84.

Roscoe, T.T., Guilleminot, J., Bessoule, J.-J., Berger, F., and Devic, M. (2015). Complementation of Seed Maturation Phenotypes by Ectopic Expression of ABSCISIC ACID INSENSITIVE3, FUSCA3 and LEAFY COTYLEDON2 in Arabidopsis. Plant Cell Physiol. 56: 1215–1228.

Santos-Mendoza, M., Dubreucq, B., Baud, S., Parcy, F., Caboche, M., and Lepiniec, L. (2008a). Deciphering gene regulatory networks that control seed development and maturation in Arabidopsis. Plant J. 54: 608–620.

Santos-Mendoza, M., Dubreucq, B., Baud, S., Parcy, F., Caboche, M., and Lepiniec, L. (2008b). Deciphering gene regulatory networks that control seed development and maturation in Arabidopsis. Plant J. 54: 608–620.

Santos Mendoza, M., Dubreucq, B., Miquel, M., Caboche, M., and Lepiniec, L. (2005). LEAFY COTYLEDON 2 activation is sufficient to trigger the accumulation of oil and seed specific mRNAs in Arabidopsis leaves. FEBS Lett. 579: 4666–4670.

Schneider, C.A., Rasband, W.S., and Eliceiri, K.W. (2012). NIH Image to ImageJ: 25 years of image analysis. Nat. Methods 9: 671–675.

Shi, L., Katavic, V., Yu, Y., Kunst, L., and Haughn, G. (2012). Arabidopsis glabra2 mutant seeds deficient in mucilage biosynthesis produce more oil. Plant J. 69: 37–46.

Singh, R. et al. (2013). Oil palm genome sequence reveals divergence of interfertile species in Old and New worlds. Nature 500: 335–339.

Song, S.-K., Ryu, K.H., Kang, Y.H., Song, J.H., Cho, Y.-H., Yoo, S.-D., Schiefelbein, J., and Lee, M.M. (2011). Cell fate in the Arabidopsis root epidermis is determined by competition between WEREWOLF and CAPRICE. Plant Physiol. 157: 1196–1208.

Stephenson, P., Baker, D., Girin, T., Perez, A., Amoah, S., King, G.J., and Østergaard, L. (2010). A rich TILLING resource for studying gene function in Brassica rapa. BMC Plant Biol. 10: 62.

Stone, S.L., Kwong, L.W., Yee, K.M., Pelletier, J., Lepiniec, L., Fischer, R.L., Goldberg, R.B., and Harada, J.J. (2001). LEAFY COTYLEDON2 encodes a B3 domain transcription factor that induces embryo development. Proc. Natl. Acad. Sci. U. S. A. 98: 11806–11811.

Tsai, A.Y.-L. and Gazzarrini, S. (2012). AKIN10 and FUSCA3 interact to control lateral organ development and phase transitions in Arabidopsis. Plant J. 69: 809–821.

Wang, J., Liu, H., and Ren, G. (2014). Near-infrared spectroscopy (NIRS) evaluation and regional analysis of Chinese faba bean (Vicia faba L.). The Crop Journal 2: 28–37.

Wang, S. et al. (2015). The OsSPL16-GW7 regulatory module determines grain shape and simultaneously improves rice yield and grain quality. Nat. Genet. 47: 949–954.

Willmann, M.R., Mehalick, A.J., Packer, R.L., and Jenik, P.D. (2011). MicroRNAs regulate the timing of embryo maturation in Arabidopsis. Plant Physiol. 155: 1871–1884.

Wu, F.-H., Shen, S.-C., Lee, L.-Y., Lee, S.-H., Chan, M.-T., and Lin, C.-S. (2009). Tape-Arabidopsis Sandwich-a simpler Arabidopsis protoplast isolation method. Plant Methods 5: 16.

Yau, R. and Rape, M. (2016). The increasing complexity of the ubiquitin code. Nat. Cell Biol. 18: 579–586.

Zhai, Z., Liu, H., and Shanklin, J. (2017). Phosphorylation of WRINKLED1 by KIN10 Results in Its Proteasomal Degradation, Providing a Link between Energy Homeostasis and Lipid Biosynthesis. Plant Cell 29: 871–889.

Zhang, X., Garreton, V., and Chua, N.-H. (2005). The AIP2 E3 ligase acts as a novel negative regulator of ABA signaling by promoting ABI3 degradation. Genes Dev. 19: 1532–1543.

Zhou, Y. et al. (2013). HISTONE DEACETYLASE19 interacts with HSL1 and participates in the repression of seed maturation genes in Arabidopsis seedlings. Plant Cell 25: 134–148.

